# FAM134B regulates ER-to-lysosome-associated degradation of misfolded proteins upon pharmacologic or genetic inactivation of ER-associated degradation

**DOI:** 10.1101/2023.11.28.569025

**Authors:** Elisa Fasana, Ilaria Fregno, Carmela Galli, Maurizio Molinari

**Affiliations:** Faculty of Biomedical Sciences, Institute for Research in Biomedicine, Università della Svizzera italiana (USI), 6500 Bellinzona, Switzerland; School of Life Sciences, École Polytechnique Fédérale de Lausanne, 1015 Lausanne, Switzerland

**Keywords:** Endoplasmic reticulum (ER), ER-associated degradation (ERAD), ER-to-lysosome-associated degradation (ERLAD), ER-phagy, Endolysosome, Proteasome, Protein quality control

## Abstract

About 40% of the eukaryotic cell’s proteome is synthesized and assembled in the endoplasmic reticulum (ER). Native proteins are transported to their intra- or extra-cellular site of activity. Folding-defective polypeptides are dislocated across the ER membrane into the cytoplasm, poly-ubiquitylated and degraded by 26S proteasomes (ER-associated degradation, ERAD). Large misfolded proteins like mutant forms of collagen or aggregation-prone mutant forms of alpha1 antitrypsin cannot be dislocated across the ER membrane for ERAD. Rather, they are segregated in ER subdomains that vesiculate and deliver their cargo to endolysosomal compartments for clearance by ER-to-lysosome-associated degradation (ERLAD). Here, we show the lysosomal delivery of a canonical ERAD substrate upon pharmacologic and genetic inhibition of the ERAD pathways. This highlights the surrogate intervention of ERLAD to remove defective gene products upon dysfunctional ERAD.

## Introduction

Maintenance of cellular homeostasis relies on efficient clearance of defective gene products. The ER is site of gene expression in nucleated cells and hosts an efficient quality control machinery that distinguishes native proteins to be delivered at their site of activity (i.e., organelles of the endocytic and secretory pathways, intracellular and plasma membranes, extracellular space), from terminally misfolded polypeptides to be removed from cells. Dislocation of misfolded proteins into the cytosol for proteasomal clearance (ERAD) is a major pathway that operates in eukaryotic cells to remove folding-defective proteins generated in the ER lumen and membranes (**Fig. 1A**) (Christianson *et al*, 2023; Sun & Brodsky, 2019). Recently, it has been observed that large misfolded proteins fail retro-translocation into the cytosol for proteasomal degradation. These polypeptides are segregated in specialized ER subdomains that separate from the biosynthetic organelle and are delivered to degradative endolysosomal/vacuolar compartments for clearance by mechanistically distinct ER-phagy pathways collectively defined as ER-to-lysosome-associated degradation ERLAD (**Fig. 1B**) (Rudinskiy & Molinari, 2023).

**Figure 1.**
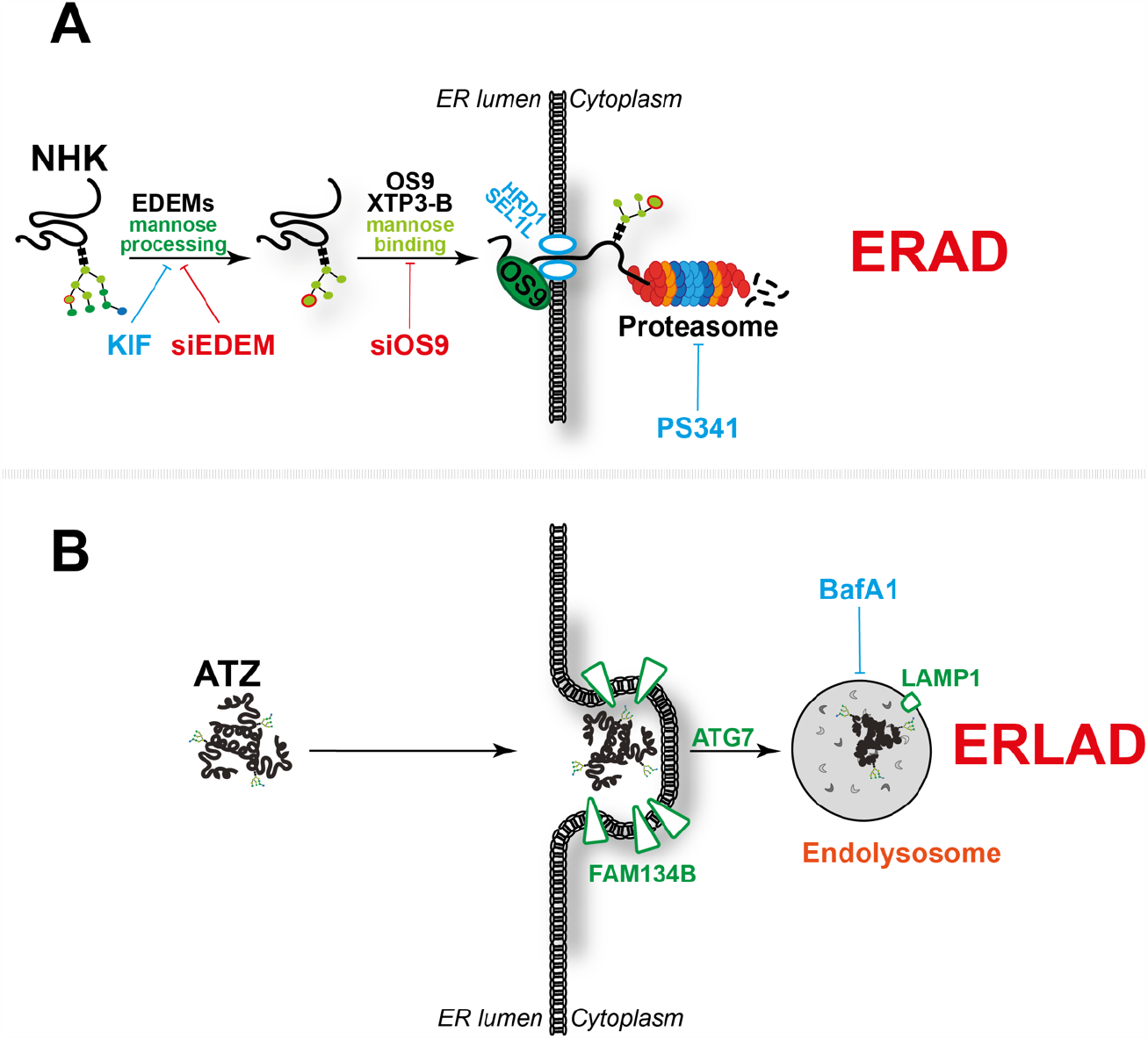
Schematic representation of proteasomal and lysosomal clearance of misfolded proteins from the ER. **A** ER-associated degradation (ERAD) pathways rely on mannose processing that generates a terminal a1,6 linked mannose residue (red circle) that engages mannose-binding lectins to facilitate dislocation of misfolded proteins across the ER membrane for proteasomal degradation. **B** ER-to-lysosome-associated (ERLAD) pathways rely on segregation of misfolded proteins in ER subdomains displaying ER-phagy receptors at the limiting membrane and their delivery to degradative compartments.

The NHK variant of α1 antitrypsin is a disease-causing folding-defective glycoprotein (Sifers *et al*, 1988). It is probably the best-characterized client of the ERAD machinery. Clearance of NHK is substantially delayed upon inhibition of de-mannosylation of NHK-bound N-glycans or upon inactivation of cytosolic 26S proteasomes (Liu *et al*, 1997, 1999). N-glycans de-mannosylation interrupts futile folding attempts in the ER lumen and engages OS9 ERAD lectins and a retro-translocation machinery including SEL1L and the E3 ubiquitin ligase HRD1 (Bernasconi *et al*, 2010; Bernasconi *et al*, 2008; Chiritoiu *et al*, 2020; Christianson *et al*, 2008; Hirao *et al*, 2006; Molinari *et al*, 2003; Oda *et al*, 2003; Olivari *et al*, 2006). KIF selectively inactivates the members of the glycosyl hydrolase 47 family of α1,2-mannosidases (which includes the α1,2-mannosidase I and the three EDEM proteins) (KIF, **Fig. 1A**) (Moremen & Molinari, 2006; Olivari & Molinari, 2007) and substantially delays ERAD of NHK (Liu *et al*., 1999; Vallee *et al*, 2000). NHK is dislocated into the cytosol, poly-ubiquitylated and degraded by 26S proteasomes, whose inhibition with the dipeptide boronic acid Bortezomib/PS341 (Adams *et al*, 1999) also prevents NHK dislocation across the ER membrane (**Fig. 1A**).

The Z-variant of α1 antitrypsin (ATZ) forms large polymers that do not enter the ERAD pathways. Rather, ATZ polymers are degraded by autophagic pathways (Chu *et al*, 2014; Hidvegi *et al*, 2010; Kroeger *et al*, 2009; Pastore *et al*, 2013; Teckman & Perlmutter, 2000) upon segregation in ER subdomains decorated with the ER-phagy receptor FAM134B that are eventually delivered to endolysosomes for clearance (**Fig. 1B**) (Fregno *et al*, 2018; Fregno *et al*, 2021). As such, inactivation of lysosomal hydrolases with bafilomycin A1 (BafA1) (Klionsky *et al*, 2008) substantially delays ATZ clearance, which accumulates within LAMP1-positive endolysosomes (Fregno *et al*., 2018; Fregno *et al*., 2021).

Notably, ERAD inhibition delays, rather than blocking degradation of ERAD clients, hinting at alternative pathways intervening to ensure efficient clearance of misfolded proteins generated in the ER under condition of ERAD impairment or overload (Molinari, 2007). Here, we monitor the fate of the canonical ERAD client NHK under conditions that perturb engagement of the ERAD machinery. We report on the enhanced NHK delivery to the endolysosomal compartment to compensate pharmacologically- or genetically-induced dysfunction of the ubiquitin/proteasome system. Our data reveal that FAM134-driven ERLAD pathways do compensate ERAD loss.

## Results

### The ERAD client NHK is normally not delivered to endolysosomes for clearance

To confirm delivery of the canonical ERLAD client ATZ to LAMP1-positive endolysosomes, mouse 3T3 cells expressing HA-tagged ATZ were incubated with 50nM BafA1 for 12h to inhibit lysosomal hydrolases, thus preserving delivered material in the lumen of the organelle (Klionsky *et al*., 2008). Confocal laser scanning microscopy (CLSM) shows the delivery of ATZ in endolysosomes that display LAMP1 at the limiting membrane (**Fig. 2A** and Inset), which has been quantified by LysoQuant, a deep-learning-based analysis software for segmentation and classification of fluorescence images (**Fig. 2E**, ATZ+BafA1) (Morone *et al*, 2020). Under the same experimental conditions, the ERAD client NHK is delivered to a much lower extent within LAMP1-positive degradative compartments (**Figs. 2B, 2E**, NHK+BafA1). This is expected because NHK is an ERAD client and cell exposure to inhibitors of lysosomal activity has no impact on its cellular clearance (Liu *et al*., 1999).

**Figure 2.**
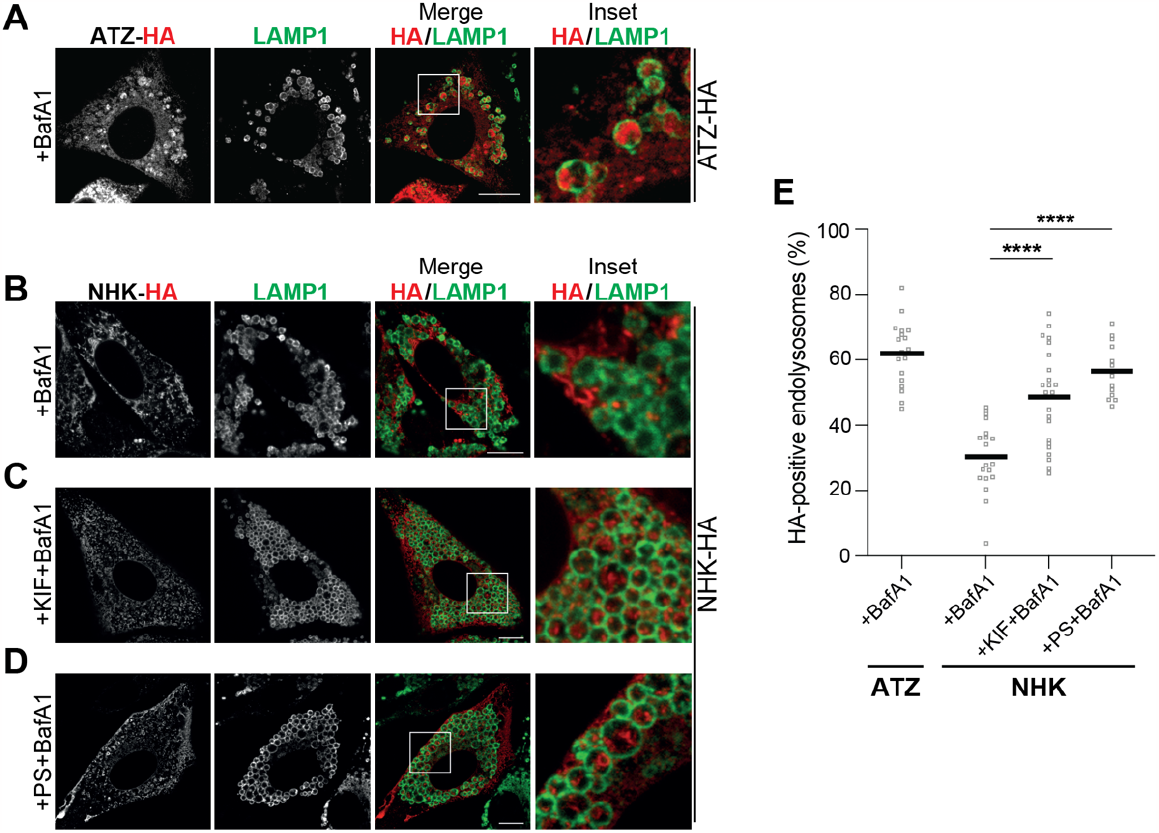
The ERAD client NHK is re-directed to endolysosomes upon pharmacologic inactivation of ERAD. **A** Clients of the ERLAD pathways (e.g., ATZ) accumulate within LAMP1-positive endolysosomes upon 3T3 cell exposure to BafA1. **B** ERAD clients such as NHK are cleared by cytosolic 26S proteasomes. They are not delivered, and do not accumulate, within LAMP1-positive endolysosomes in 3T3 cells. **C** NHK is delivered and accumulates within endolysosomes inactivated with BafA1, upon ERAD inhibition with KIF. **D** Same as **C**, when ERAD is inhibited with PS341. **E** LysoQuant quantification of ATZ (panel **A**) and NHK accumulation (panels **B-D**) within LAMP1-positive endolysosomes (n=19, 19, 23 and 13 cells, respectively). One-way analysis of variance (ANOVA) and Dunnett’s multiple comparison test, ****P<0.0001. Scale bar: 10μM.

### Pharmacologic inhibition of ERAD triggers lysosomal delivery of NHK

Next, we examined the delivery to endolysosomes of NHK upon pharmacologic inhibition of the ERAD machinery. KIF inhibits NHK clearance via ERAD (Liu *et al*., 1999), and it does so by preventing the generation of the low-mannose signal decoded by OS9 proteins that should shuttle terminally misfolded polypeptides to the SEL1L/HRD1 dislocation machinery (**Fig. 1A**) (Bernasconi *et al*., 2010; Bernasconi *et al*., 2008; Christianson *et al*., 2008). To verify if ERAD inactivation upon inhibition of α1,2-mannosidases resulted in NHK delivery to the lysosomal district, mouse 3T3 cells expressing HA-tagged NHK were incubated with 50μg/ml KIF to inhibit ERAD and with 50nM BafA1 for 12h to preserve NHK possibly delivered in the lumen of endolysosomes. CLSM reveals the enhanced accumulation of NHK in endolysosomes that display LAMP1 at the limiting membrane (**Fig. 2C**), which has been quantified by LysoQuant (**Fig. 2E**).

The second test was done by pharmacologic inhibition of the 26S proteasome with PS341(Adams *et al*., 1999), which blocks NHK clearance via ERAD (Liu *et al*., 1999), and it does so by preventing the NHK dislocation across the ER membrane, which is coupled with proteasomal activity. To this end, mouse 3T3 cells expressing HA-tagged ATZ were incubated with 10nM PS341 to inhibit ERAD and with 50nM BafA1 for 12h to preserve NHK possibly delivered in the lumen of endolysosomes. CLSM reveals the enhanced delivery of NHK in endolysosomes that display LAMP1 at the limiting membrane (**Fig. 2D**), which has been quantified by LysoQuant (**Fig. 2E**). Thus, inhibition of ERAD at early and late steps of client’s selection, activates channeling of misfolded polypeptides into ERLAD pathways that deliver them to lysosomal compartments.

### Genetic inhibition of ERAD upon silencing of EDEM1 triggers lysosomal delivery of NHK

EDEM1 is an active α1,2-mannosidase that accelerates extraction of misfolded glycoproteins from the calnexin chaperone system to direct them for proteasomal degradation (Molinari *et al*., 2003; Oda *et al*., 2003; Olivari *et al*., 2006). Silencing of EDEM1 expression substantially delays ERAD of glycoproteins (Molinari *et al*., 2003). To verify whether silencing of EDEM1 expression diverts NHK into the ERLAD pathway, we compared delivery of HA-tagged NHK in the LAMP1-positive endolysosomes in HEK293 cells transfected with a negative control (small interfering scrambled RNA, shCTRL), or with a small interfering RNA targeting the EDEM1 sequence (shEDEM, **Fig. 3A**). Cells were exposed to 50nM BafA1 for 12h to preserve the NHK fraction possibly delivered in the lumen of endolysosomes. CLSM reveals the enhanced accumulation of NHK in endolysosomes that display LAMP1 at the limiting membrane in cells with reduced EDEM1 levels (**Fig. 3B**, lower panel and Inset), which has been quantified by LysoQuant (**Fig. 3C**).

**Figure 3.**
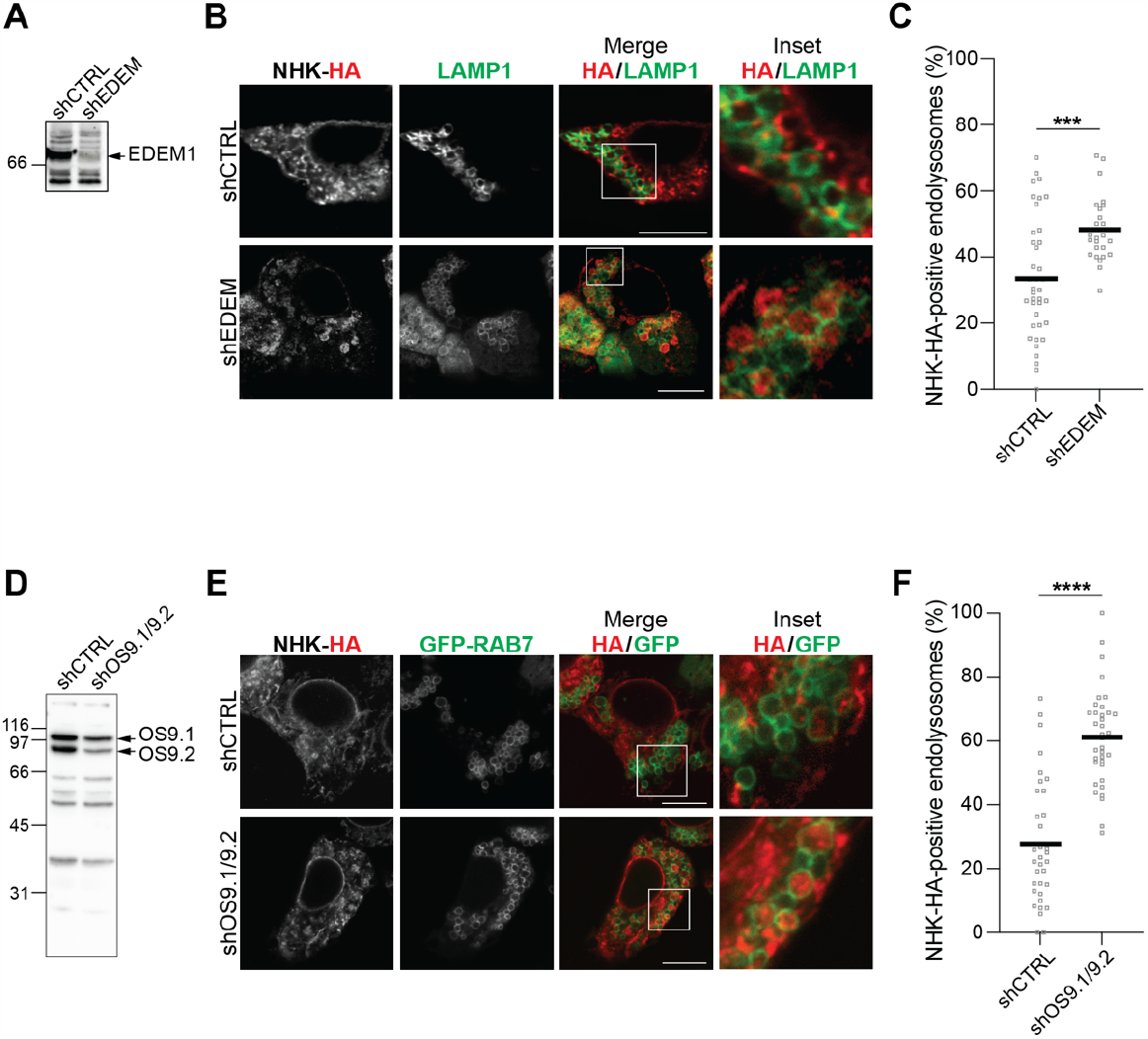
The ERAD client NHK is re-directed to endolysosomes upon genetic inactivation of ERAD. **A** Western blot showing the efficiency of EDEM1 silencing in HEK293 cells. **B** Silencing EDEM1 expression enhances NHK delivery within endolysosomes in HEK293 cells. **C** LysoQuant quantification of NHK accumulation within LAMP1-positive endolysosomes in cells with normal (shCTRL) and reduced EDEM1 levels (shEDEM) (n=39 and 25 cells, respectively). Unpaired t-test, *** P<0.001. **D** Same as **A** for OS9.1/OS9.2. **E** Silencing OS9.1 and OS9.2 expression enhances NHK delivery within endolysosomes in HEK293 cells. **F** LysoQuant quantification of NHK accumulation within LAMP1-positive endolysosomes in cells with normal (shCTRL) and reduced OS9.1 and OS9.2 levels (shOS9.1/OS9.2) (n=35 and 36 cells, respectively). Unpaired t-test, **** P<0.0001. Scale bar: 10μM.

### Genetic inhibition of ERAD upon silencing of OS9 proteins triggers lysosomal delivery of NHK

De-mannosylation of N-glycans on misfolded glycoproteins generates binding sites for the OS9 ERAD lectins, which are expressed in two ER stress-induced splice variants in the ER lumen (Bernasconi *et al*., 2008). OS-9 association facilitates proteasomal clearance of NHK by promoting delivery of the misfolded polypeptide to the HRD1/SEL1L retro-translocation machinery (**Fig. 1A**) (Bernasconi *et al*., 2010; Bernasconi *et al*., 2008; Christianson *et al*., 2008). We and others previously showed that silencing of OS9 expression substantially impairs ERAD of NHK (Bernasconi *et al*., 2008; Christianson *et al*., 2008). To verify whether under these conditions NHK was diverted into the ERLAD pathways, we monitored NHK delivery in the LAMP1-positive endolysosomes in HEK293 cells transfected with a control small interfering scrambled RNA (shCTRL), or with a small interfering RNA targeting the OS9.1 and OS9.2 sequences (shOS9.1/OS9.2, **Fig. 3D**). CLSM reveals the enhanced delivery of NHK in LAMP1-positive endolysosomes in cells where ERAD is defective upon silencing of OS9 proteins expression (**Figs. 3E** and Inset, **3F**).

### The ER-phagy receptor FAM134B drives lysosomal delivery of NHK upon ERAD inactivation

Misfolded polypeptides that fail engagement of the ERAD machinery (ATZ polymers are shown as example in **Figs. 1B** and **2A**) are segregated in ER subdomains displaying the ER-phagy receptor FAM134B at the limiting membrane and are eventually delivered to LAMP1-positive endolysosomal compartments for clearance (Fregno *et al*., 2018; Rudinskiy & Molinari, 2023). If ERAD clients would co-opt the same pathway for lysosomal clearance upon ERAD inactivation, it is expected that NHK delivery to the LAMP1-positive compartment is substantially inhibited in cells lacking FAM134B. To test this hypothesis, the experiments described in the previous sections were reproduced in mouse embryonic fibroblasts (MEF) subjected to CRISPR/Cas9 genome editing to silence the expression of FAM134B (**Fig. 4A**). NHK does not accumulate in LAMP1-positive endolysosomes in wild type MEF (**Figs. 4B**, upper panels, **4C**) and in cells lacking the ER-phagy receptor FAM134B (**Figs. 4B**, lower panels, **4C**). Inactivation of the ERAD pathway in wild type MEF upon cell exposure to KIF, promotes NHK delivery to the LAMP1-positive endolysosomal compartments (**Figs. 4D, 4K**). Delivery of NHK in the LAMP1-positive compartment is virtually abolished upon deletion of FAM134B (**Figs. 4E, 4K**, pCDNA3). The back-transfection of V5-tagged FAM134B restores NHK delivery to the LAMP1-positive endolysosomes (**Figs. 4F, 4K**). The back-transfection of a variant of FAM134B that carries mutations in the LIR domain that prevent engagement of cytosolic LC3 molecules (Fregno *et al*., 2018), is not restoring NHK delivery to endolysosomes (**Figs. 4G, 4K**). The FAM134B-driven deviation of NHK into the ERLAD pathways ensuring lysosomal clearance of misfolded proteins generated in the ER is also induced when access of NHK to the ERAD pathways is abolished upon MEF exposure to the proteasomal inhibitor PS341 (**Figs. 4H-4K**).

**Figure 4.**
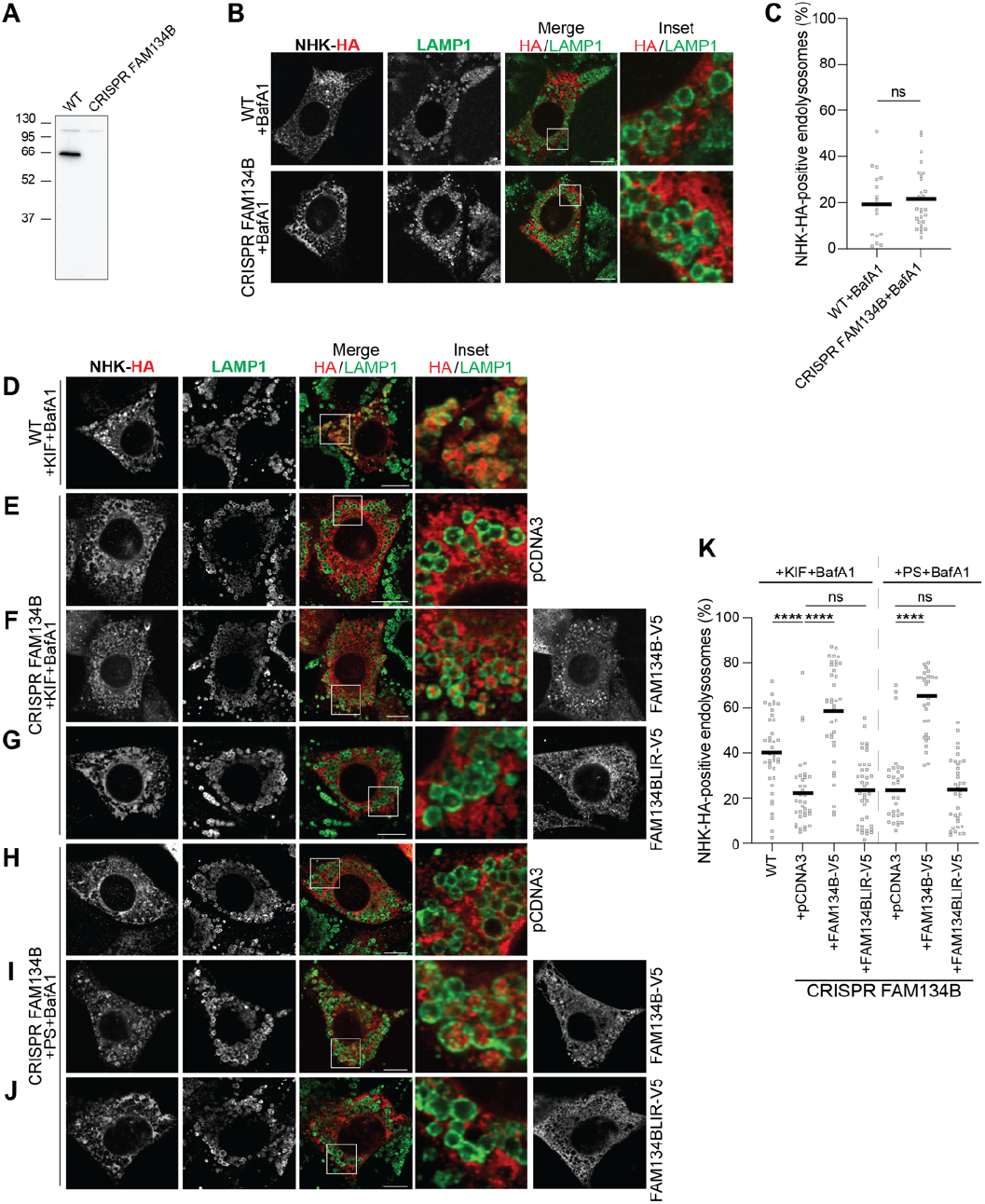
The ER-phagy receptor FAM134B drives delivery of the ERAD client NHK into the ERLAD pathway. **A** Western blot showing deletion of FAM134B in MEF. **B** NHK is an ERAD client and does not accumulate in LAMP1-positive endolysosomes in wild type (upper panels) and FAM134B-KO cells (lower panels). **C** LysoQuant quantification of **B** (n=17 and 27 cells, respectively). Unpaired t-test, ^ns^P> 0.05. **D** NHK is delivered in LAMP1-positive endolysosomes upon ERAD inhibition with KIF in wild type MEF. **E** NHK is not delivered in LAMP1-positive endolysosomes upon ERAD inhibition with KIF in MEF lacking FAM134B. **F** Back-transfection of FAM134B in FAM134B-KO MEF restores NHK delivery within endolysosomes. **G** Back-transfection of mutant FAM134B that cannot engage LC3 fails to restore NHK delivery within endolysosomes. **H** Same as **D**, where ERAD is inhibited with PS341. **I** Same as **E**, where ERAD is inhibited with PS341. **J** Same as **F**, where ERAD is inhibited with PS341. **K** LysoQuant quantification of **D-J** (n=39, 35, 37, 39, 31, 30 and 35 cells, respectively). One-way analysis of variance (ANOVA) and Dunnett’s multiple comparison test, ^ns^P> 0.05, ****P<0.0001. Scale bar: 10μM.

### Involvement of autophagy genes in compensatory ERLAD pathways

ER-phagy receptors control clearance of ER subdomains upon engagement of various autophagy gene products that will determine if clearance of ER subdomains will proceed via *macro*-ER-phagy involving double membrane autophagosomes, or via other types of ER-phagy that will not involve autophagosomes and their biogenesis (Chino & Mizushima, 2023; Reggiori & Molinari, 2022). Previous work in our lab showed that FAM134B-controlled delivery of ATZ polymers to LAMP1-positive endolysosomes involves the LC3 lipidation machinery but does not involve the autophagosome biogenesis machinery. Consistently, deletion of the autophagy gene Atg7 that abolishes LC3 lipidation (Komatsu *et al*, 2005) prevents ATZ delivery to endolysosomal compartments. In contrast, deletion of ATG13, a crucial component of the autophagosome biogenesis machinery (Hosokawa *et al*, 2009; Kaizuka & Mizushima, 2016; Suzuki *et al*, 2014) does not impact on ATZ delivery to the degradative district (Fregno *et al*., 2018). The repetition of these experiments to monitor the case of the ERAD client NHK confirms that the misfolded protein is not normally delivered to the LAMP1-positive endolysosomal compartment of MEF (**Figs. 5A, 5H** and **Fig. 2**), unless the ERAD pathway has been inactivated upon cell exposure to the α1,2-mannosidases inhibitor KIF or the proteasomal inhibitor PS341 (**Figs. 5B, 5C, 5H** and **Fig. 2**). In cells lacking ATG7 (**Fig. 5D**), the delivery of NHK to the LAMP1-positive degradative compartment is abolished (**Figs. 5E, 5K**). In cells lacking ATG13 (**Fig. 5F**), the delivery of NHK to the LAMP1-positive degradative compartment proceeds unperturbed (**Figs. 5G, 5H**). These results recapitulate the phenotype previously observed for clearance of ATZ polymers, which is hampered in cells with defective LC3 lipidation (Chu *et al*., 2014; Fregno *et al*., 2018; Hidvegi *et al*., 2010; Kroeger *et al*., 2009; Pastore *et al*., 2013; Teckman & Perlmutter, 2000), but remains unaffected in cells with defective autophagosome biogenesis (Fregno *et al*., 2018). All in all, perturbation of the ERAD pathway deviates the ERAD client NHK into the catabolic ERLAD pathway normally used to remove ATZ polymers from cells.

**Figure 5.**
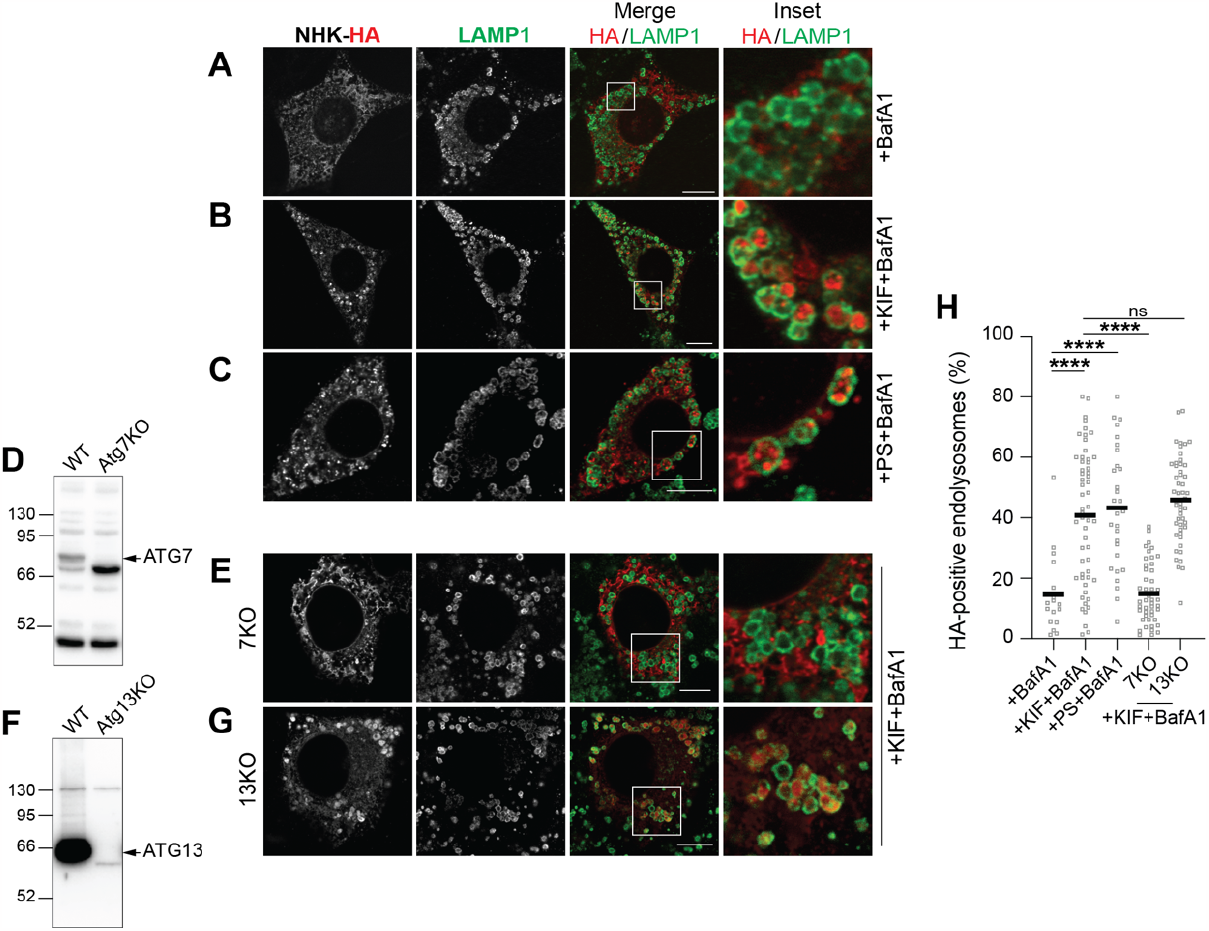
The LC3 lipidation is involved, autophagosome biogenesis is dispensable, for delivery of NHK to ERLAD upon perturbation of the ERAD machinery. **A-C** Same as **Figs. 2B-2D**, in MEF. **D** Western blot showing deletion of ATG7 in MEF. **E** NHK does not accumulate in LAMP1-positive endolysosomes in ATG7-KO cells. **F** Western blot showing deletion of ATG13 in MEF. **G** NHK delivery within LAMP1-positive endolysosomes is not perturbed in ATG13-KO MEF. **H** LysoQuant quantification of **A-C, E, G** (n=17, 57, 29, 46 and 50 cells, respectively). One-way analysis of variance (ANOVA) and Dunnett’s multiple comparison test, ^ns^P> 0.05, ****P<0.0001. Scale bar: 10μM.

## Discussion

A stringent protein quality control operates in the ER of eukaryotic cells. Proteins that have completed their folding program are released from ER-resident molecular chaperones, exit the ER to be delivered at their final intra- or extra-cellular destination. Folding is error prone, and the rate of misfolding is substantially enhanced by mutations in the polypeptide sequence. The incapacity to fold correctly, results in selection of the aberrant gene product for degradation, or the formation of aggregates that are retained in the biosynthetic compartment. Dedicated machineries are available in the ER lumen and membranes to distinguish misfolded or incompletely folded polypeptides to be retained in the ER lumen, from native and functional proteins to be released. Incompletely folded polypeptides retained in the ER are exposed to the folding environment and can eventually reach the native, transport-permissive architecture. When folding is impossible, the polypeptides are actively extracted from the ER, are translocated across the ER membrane and are degraded by proteasomes in pathways named ERAD (**Fig. 1A**). An increasing number of misfolded proteins is emerging in the literature that cannot enter ERAD pathways (Rudinskiy & Molinari, 2023). In many cases, these are large polypeptides or polypeptides that are prone to form aggregates or polymers. A variety of ERLAD pathways are available in nucleated cells that sequester these ERAD-resistant polypeptides in dedicated subdomains that are eventually separated from the ER and form ER-derived vesicles that are either engulfed by autophagosomes, by endolysosomes, or fuse with degradative compartments releasing their cargo in the hydrolytic lumen (**Fig. 1B**). Our study shows that the intervention of ERLAD pathways is not limited to clearance of large proteins that fail to be dislocated across the ER membrane for ERAD. Rather, ERLAD may be engaged by ERAD clients when their preferred road-to-destruction is dysfunctional or saturated by an excess of aberrant gene products (**Fig. 6**).

**Figure 6.**
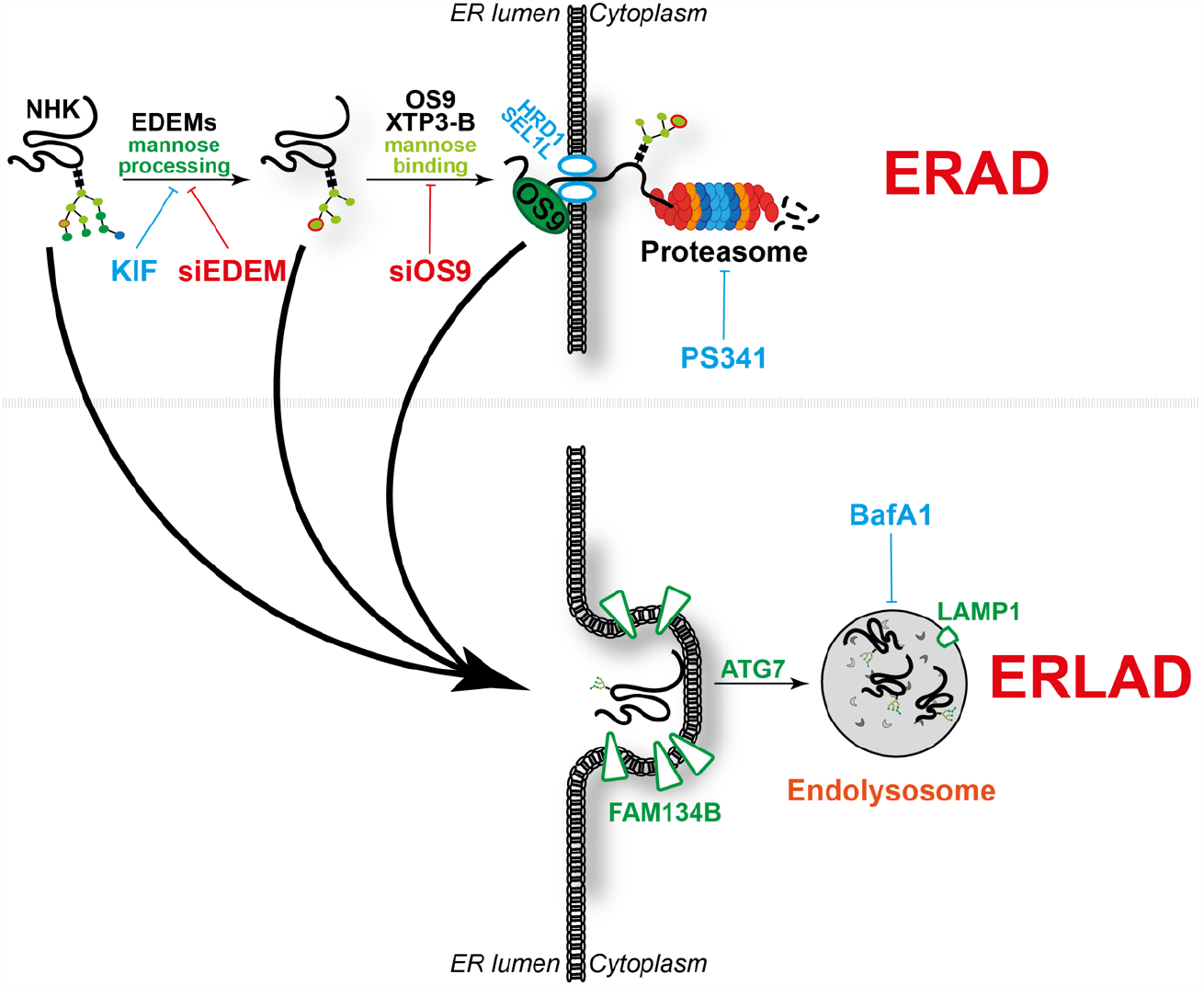
Schematic representation of how pharmacologic and genetic ERAD perturbation triggers compensatory ERLAD.

## Methods

### Expression plasmids and antibodies

HA-tagged ATZ and NHK and V5-tagged FAM134B and FAM134BLIR are subcloned in pcDNA3.1 plasmids. pDest-EGFP-Rab7 is a kind gift from T. Johansen. The polyclonal anti-HA (Sigma cat. H6908), the anti-V5 (Invitrogen cat. 46-0705) and the anti-LAMP1 (Hybridoma Bank, 1D4B deposited by J.T. August) are described in (Fregno *et al*., 2021). The anti-FAM134B is a kind gift of M. Miyazaki (University of Colorado Denver); the anti-EDEM1 is from Sigma (cat. E8406); the anti OS-9 is from Novus Biologicals (cat. BC100-520); the anti-ATG7 and anti-ATG13 are from Sigma (cat. SAB4200304 and SAB4200100, respectively). Alexa-conjugated secondary antibodies are from Thermofisher; HRP-conjugated Protein A is from Invitrogen; HRP-conjugated anti-mouse is from SouthernBiotech).

### Cell Lines, transient transfections, pharmacologic inhibition

Flp-InTM-3T3 cells (Thermo Fisher) stably expressing Halo-ATZ-HA or Halo-NHK-HA were generated following manufacturer instructions and cultured in DMEM supplemented with 10% FCS and 150μg/ml Hygromycin. MEF and HEK293 cells were grown in DMEM/10% FBS. HEK293FT cells expressing reduced levels of OS9.1 and OS9.2 are described in (Bernasconi *et al*., 2008). FAM134B-deficient MEF cells (CRISPR FAM134B) were generated using CRISPR/Cas9 genome editing protocol as described in (Fumagalli *et al*, 2016).

Transient transfections were performed using JetPrime transfection reagent (PolyPlus) following manufacturer’s protocol. BafA1 (Calbiochem) was used at 50 nM for 12 h; KIF (Toronto Research Chemicals) was used at 200μM for 12h; PS341 (LubioScience) was used 12h at 100nM for Flp-InTM-3T3 cells or at 5nM for MEF cells.

### Confocal laser scanning microscopy

Cells plated on Alcian Blue-treated glass coverslips are washed with PBS and fixed at room temperature for 20 min in 3.7% formaldehyde diluted in PBS. Cells are permeabilized for 15 min with 0.05% saponin, 10% goat serum, 10mM HEPES, 15mM glycine (PS) and exposed for 90 min to primary antibodies diluted 1:100 in PS. After washes with PS, cells are incubated for 45 min with Alexa Fluor-conjugated secondary antibodies diluted 1:300 in PS. Cells are rinsed with PS and water and mounted with Vectashield (Vector Laboratories). Confocal images are acquired on a Leica TCS SP5 microscope with a Leica HCX PL APO lambda blue 63.0X1.40 OIL UV objective. Quantitative analyses are performed with LysoQuant as described in (Morone *et al*., 2020).

### Statistical analysis

Plots and statistical analyses were performed using GraphPad Prism10 (GraphPad Software Inc.). In this study, one-way ANOVA with Dunnett’s multiple comparisons test and unpaired t-test were used to assess statistical significance. An adjusted P-value < 0.05 was considered as statistically significant.

## Notes

### Competing Interest Statement

The authors have declared no competing interest.

